# Tobacco TGA7 mediates gene expression dependent and independent of salicylic acid

**DOI:** 10.1101/341834

**Authors:** Vlatka Stos-Zweifel, David Neeley, Evelyn Konopka, Meike Meissner, Meike Hermann, Felix Maier, Verena Häfner, Artur J.P. Pfitzner, Ursula M. Pfitzner

## Abstract

Basic region leucine zipper (bZIP) transcription factors of the TGA family control gene expression in response to diverse stimuli. *Arabidopsis* clade II and clade III TGA factors mediate salicylic acid (SA)-induced expression of *PATHOGENESIS-RELATED GENE1* (*PR-1*) via interplay with NONEXPRESSOR OF PR GENES1 (NPR1, a.k.a. NIM1). Interaction with TGA factors occurs through the central ankyrin repeat domain of NPR1. In a yeast two-hybrid screen with the NPR1 bait, we identified TGA7, a novel member of the tobacco (Nt) TGA family grouping to clade III. TGA7 is most similar to NtTGA1a, and, like NtTGA1a, TGA7 displays transcription activity in yeast. Unexpectedly, TGA7 preferentially and uniquely interacts with the SA-sensitive C-terminal region of NtNPR1, demonstrating that NtNPR1 harbors multiple distinct TGA factor binding sites. Interaction with NPR1 impairs TGA7 transcription activity in yeast. Furthermore, TGA7 binding to the NtNPR1 C-terminus is outcompeted by SA-induced type 2 NIM1-INTERACTING (NIMIN) proteins. In tobacco plants, a TGA7–Gal4 DNA-binding domain chimeric protein (TGA7GBD) mediates SA-responsive reporter gene expression in young leaf tissue and spontaneous reporter activation in older leaves displaying *PR-1* gene expression. Astonishingly, TGA7GBD is also able to activate the reporter independent from *PR-1* gene expression in noninduced cotyledons of tobacco seedlings. Together, our findings support a model in which TGA7 mediates both SA-dependent and SA-independent gene activation controlled by the plant’s developmental stage and by the C-terminal region of constitutively accumulating NtNPR1.

## INTRODUCTION

Response of cells to external and internal signals requires reprograming mediated by transcription factors which bind to specific sites in the promoter regions thus controlling expression of their target genes. In plants, many transcription factors are organized in large multigene families whose members may play roles in different processes, like growth, development and response to biotic and abiotic stress. A potent group of transcription factors found in eukaryotes are basic (b) region leucine zipper (ZIP) motif proteins. bZIP transcription factors bind to DNA through their basic region and dimerize through the hydrophobic sides of their heptad repeat of leucines (Landschulz et al., 1988; Ellenberger et al., 1992). Plant bZIP factors regulate diverse biological processes including pathogen defense, light and stress signaling, seed maturation and flower development (Jakoby et al., 2002). TGA transcription factors are a subgroup of plant bZIP proteins (Jakoby et al., 2002; Gatz, 2013). They bind to cis-regulatory element *activating sequence-1* (*as-1*), originally identified in the *Cauliflower mosaic virus* (*CaMV*) *35S* promoter, encompassing a tandem repeat of the sequence TGACG (Lam et al., 1989; Katagiri et al., 1989).

In *Arabidopsis*, the TGA family comprises ten members, grouped in five clades (Jakoby et al., 2002). Proteins of clades IV (TGA9, TGA10) and V (PERIANTHIA, PAN) are involved in floral development (Chuang et al., 1999; Murmu et al., 2010), while factors of clades I to III have been associated with pathogen defense reactions. Initially, it was established that salicylic acid (SA)-induced *PATHOGENESIS-RELATED (PR)* genes of group 1 from tobacco and *Arabidopsis* contain SA-responsive cis-regulatory elements related to viral *as-1* (Strompen et al., 1998; Lebel et al., 1998). SA is a signal molecule in systemic acquired resistance (SAR) protecting plants against secondary pathogen infection after an initial attack causing cell death at the pathogen invasion sites, known as hypersensitive reaction (HR; Vlot et al., 2009). In vitro DNA-protein binding studies confirmed that TGA factors can access *as-1*-related elements in the tobacco (Nt) and *Arabidopsis* (At) *PR-1* promoters (Strompen et al., 1998; Zhang et al., 1999; Zhou et al., 2000; Després et al., 2000), and chromatin immunoprecipitation (ChIP) assays demonstrated that clade II and clade III factors TGA2 and TGA3 are recruited to the *AtPR-1* promoter in planta (Johnson et al., 2003; Rochon et al., 2006). Further strong support for a crucial role of members of the TGA family in pathogen defense arose after cloning the central SAR regulator *NONEXPRESSOR OF PR GENES1* (*NPR1*) from *Arabidopsis* (Cao et al., 1997; Ryals et al., 1997; Shah et al., 1997). NPR1 is indispensable for SA-dependent *PR-1* expression. Yeast two-hybrid (Y2H) screens using AtNPR1 as bait uncovered TGA factors as interaction partners of NPR1 (Zhang et al., 1999; Després et al., 2000; Zhou et al., 2000), thus establishing a physical link between the central SAR regulator and SA-induced expression of the SAR effector *PR-1*. NPR1 interactors belong to clade II (TGA2, TGA5, TGA6) and clade III (TGA3, TGA7), and it was suggested that TGA2 and TGA3 bind to the ankyrin repeat domain in the central region of NPR1, but the NPR1 N-terminus is needed to facilitate strong interaction with these factors (Zhang et al., 1999; Zhou et al., 2000).

The same TGA factors, which bind to NPR1, interact also with its paralogs, NPR3 and NPR4. NPR3 and NPR4 have been implied in pathogen defense as well (Liu et al., 2005; Zhang et al., 2006; Shi et al., 2013), and recent evidence has revealed that all, NPR1, NPR3 and NPR4, are able to perceive the SA signal, albeit with very different sensitivities (Maier et al., 2011; Fu et al., 2012; Wu et al., 2012; Manohar et al., 2015). Thus, NPR family members appear to transduce the SA signal with help of clade II and clade III TGA factors. Yet, the precise actions of TGAs in transcription complexes on defense gene promoters have remained elusive. On one side, clear evidence has been provided for a positive role of clade II factors, in particular TGA2, in SA and *NPR1*-dependent gene induction by biochemical and genetic means (Fan and Dong, 2002; Zhang et al., 2003; Rochon et al., 2006; Kesarwani et al., 2007). On the other side, overexpression or inhibition of *TGA2* had no significant effect on *PR-1* gene expression after pathogen infection or treatment with the SA analog 2,6-dichloroisonicotinic acid (INA; Kim and Delaney, 2002). Moreover, evidence for a negative role of TGA2 and other clade II factors in *PR-1* gene expression has also emerged (Zhang et al., 2003; Rochon et al., 2006; Kesarwani et al., 2007). According to a model proposed by Boyle and associates, TGA2 binds initially as an oligomer to the cognate TGACG sequence of its target promoter, thereby repressing gene transcription in untreated cells. After stimulation with SA, however, an enhanceosome is assembled on the target promoter containing NPR1 and TGA2 proteins with a stoichiometry of 2:2 (Boyle et al., 2009). Furthermore, clade II TGA factors were also reported to be required for activation of the jasmonic acid (JA)/ethylene (ET)-dependent defense pathway against necrotrophic pathogens (Zander et al., 2010). Merely, analysis of the *tga3-1* mutant has revealed that this clade III factor clearly has a prominent role as transcriptional activator of *PR-1* (Kesarwani et al., 2007).

Although clade I factors TGA1 and TGA4 do not bind to NPR1 in Y2H assays (Després et al., 2000; Zhou et al., 2000; Niggeweg et al., 2000a), it was shown that treatment with SA induces interaction between NPR1 and TGA1 in *Arabidopsis* leaves (Després et al., 2003). Further analysis revealed that two C-terminal cysteine residues in TGA1, Cys-260 and Cys-266, preclude interaction with NPR1 due to formation of an intramolecular disulfide bridge. But under reducing conditions, as encountered in SA-treated plant tissue, TGA1 can bind NPR1. The data indicate that clade I factors may also play a role in pathogen defense, albeit different from the roles of clade II and clade III factors. Accordingly, Kesarwani and associates have found that TGA1 and TGA4 positively regulate basal pathogen resistance, but these factors have only marginal effects on *PR-1* gene induction by INA (Kesarwani et al., 2007). In fact, the function of clade I TGA factors has recently been established to be substantially independent of NPR1 (Shearer et al., 2012). Taken together, TGA factors of clades I to III appear crucial in various signal transduction pathways leading to *PR* gene induction and pathogen defense, and factors of clades II and III seem to exert this function via NPR1. However, the exact molecular mechanisms how TGA factors activate defense genes are not fully understood. Furthermore, TGA factors not only bind to members of the NPR family. *Arabidopsis* TGA factors also interact with glutaredoxins (GRXs) of the ROXY class (Ndamukong et al., 2007; Li et al., 2009; Zander et al., 2012), which are marked by a CC-type active site motif specifically found in GRXs of land plants (Xing et al., 2006). Glutaredoxins control the redox state of proteins by catalyzing thiol disulfide reductions (Buchanan and Balmer, 2005).

In tobacco, five TGA factors are known, falling into three groups (Gatz, 2013). The *NtTGA1a* cDNA was the first transcription factor gene cloned from plants (Katagiri et al., 1989). NtPG13 is similar to NtTGA1a (Fromm et al., 1991). The two proteins group to clade I. cDNAs coding for NtTGA2.1 and NtTGA2.2 were isolated using *AtTGA5* as probe (Niggeweg et al., 2000a). The proteins are most closely related to *Arabidopsis* TGA2 from clade II. NtTGA2.2 homodimer has been found to be the major component of transcription complex *as-1*-binding factor-1 (ASF-1) assembling on SA-responsive promoters, while NtTGA2.1 is a minor component (Niggeweg et al., 2000b). Consistently, reduction of class II factors NtTGA2.2 and NtTGA2.1 by RNA interference (RNAi) correlates with reduced expression of *PR-1a* in tobacco (Thurow et al., 2005). Both NtTGA2.2 and NtTGA2.1 bind to *Arabidopsis* NPR1 (Niggeweg et al., 2000a). Furthermore, NtTGA2.1 and NtTGA2.2 can form heterodimers in vitro and in vivo (Kegler et al., 2004). Finally, NtTGA10, like NtTGA1a, does not interact with AtNPR1 (Schiermeyer et al., 2003). This factor defines a distinct class of its own. In an effort to understand SA signaling through tobacco NPR1, we searched a tobacco cDNA library (Börnke, 2005) using the yeast two-hybrid (Y2H) system and AtNPR1 as bait protein. In this screen, we isolated three cDNA clones encoding NIM1-INTERACTING (NIMIN) proteins of type 2 (Zwicker et al., 2007) and three cDNA clones coding for a novel member of the tobacco TGA transcription factor family, which we named TGA7. The novel protein groups to clade III. TGA7 is unique among known TGA factors in that it binds to the C-terminal region of NtNPR1 which senses the SAR signal SA. There, TGA7 competes with binding of SA-induced NIMIN proteins.

## MATERIALS AND METHODS

### Plant growth conditions and chemical treatments

Tobacco plants (*N. tabacum* cv. Samsun NN, *N. benthamiana* and various transgenic tobacco lines) used for RNA and protein extraction were grown in commercial potting soil under natural light conditions in a greenhouse. Seedlings were cultivated in a growth chamber under controlled environmental conditions (20°C, 16h light/8h dark cycle and 60% relative humidity). Seeds were sown under sterile conditions on Murashige & Skoog (MS) medium without addition of sucrose, vitamins and hormones, unless indicated otherwise. Transgenic seedlings grew in presence of appropriate antibiotics. For *PR-1* gene induction, 0.3mM salicylic acid was added to the medium, and for activation of the GVG transcription factor, the medium was supplemented with 0.01mM dexamethasone (DEX) in 3% ethanol. GUS assays were performed three to four weeks after sowing the seeds on MS medium. Chemical treatment of greenhouse-grown plants was as described previously (Zwicker et al., 2007). Leaf disks were cut from young tobacco plants and floated for 3 days on water or solutions of 1mM SA, 0.34mM BTH or 1mM 4-OH BA. Treatments with other plant hormones were at a concentration of 100µM for each substance in 0.1% ethanol. Alternatively, greenhouse-grown plants were induced by spraying with water or 5mM SA, and leaf tissue was harvested after four days for extraction of proteins or RNAs.

### Isolation of clones from a tobacco yeast two-hybrid cDNA library

A Y2H cDNA library prepared from poly(A) RNA from tobacco (*N. tabacum* cv. Samsun NN) source leaves (Börnke, 2005) was screened as described previously (Weigel et al., 2001; Zwicker et al., 2007). With GBD–AtNPR1 as bait, six clones were isolated. Three clones were derived from the same gene coding for an unknown TGA transcription factor. The longest clone, pAD#1, contained a 1.2 kb cDNA insert encompassing an open reading frame of 257 amino acids fused in-frame to the transcription activation domain of the prey plasmid. The amino acid sequence starts with the first leucine of a three-digit zipper motif. The clone also contains the sequence encoding the whole conserved C-terminus, the translation stop codon, a 392 bp-long 3’-untranslated region and a poyA tail encompassing 22 adenines. The 5’-end of the cDNA was identified on two independent clones overlapping by 254 bp with the clone from the screen, which were amplified from DNA isolated from the tobacco cDNA library with primers GADfwd and TGA-8 (Table S2). The longest clone carries a cDNA insert of 735 bp encoding an open reading frame of 190 amino acids. This clone comprises the sequence of the whole N-terminal region including the highly conserved basic DNA-binding domain and the leucine zipper motif. The first ATG of the open reading frame is preceded by 164 bp of 5’-untranslated sequence including several in-frame stop codons.

The same cDNA library was screened with the GBD–TGA7Δ1 bait lacking transcription activation potential (Figures S4A and S4B and Figure S5). In total, 29 clones were identified. By far most clones (24) encode a protein identical to GLUTAREDOXIN-C6 predicted from the genome of *N. sylvestris*. The longest clone, pAD#29-3, contains the whole open reading frame of 147 amino acids, 19 bp of the 5′-untranslated region, a 217 bp-long 3′-untranslated region and a poyA tail encompassing 18 adenines. Three clones isolated in the Y2H screen code for two other glutaredoxin proteins (NtGRXC6-like and NtGRXC9-like), one clone contains a partial *NPR1* sequence, and another clone (pAD#33) contains a partial *TGA7* sequence.

### Phylogenetic analysis

The analysis was performed on the Phylogeny.fr platform using the “One Click“software developed by Dereeper et al. (2010). Briefly, protein sequences were aligned with MUSCLE (v3.8.31). After alignment, ambiguous regions were removed with Gblocks (v0.91b). The phylogenetic tree was reconstructed using the maximum likelihood method implemented in the PhyML program (v3.1/3.0 aLRT). Graphical representation and edition of the phylogenetic tree were performed with TreeDyn (v198.3).

### Protein-protein and protein-DNA interaction assays in yeast

Protein-protein interaction assays in yeast (Y1H, Y2H and Y3H) were conducted as reported earlier (Weigel et al., 2001; Maier et al., 2011; Hermann et al., 2013). To determine binding of TGA factors to DNA, the *as-1* cis-acting element from the *CaMV 35S* promoter was inserted in reporter plasmid pRW95-3 (Wolf et al., 1996). pRW95-3/as-1 was transformed together with pGAD424/TGA plasmids in yeast. In all Y1H, Y2H and Y3H analyses, *lacZ* reporter gene activities were tested in duplicate with at least three independent colonies. The experiments were repeated at least once with new yeast transformations. Representative results collected in one experimental set are shown. The results are depicted as mean enzyme activities in Miller units, plus and minus standard deviation (SD).

### Detection of GFP fusion proteins in yeast and *Nicotiana benthamiana*

Expression constructs pEGFP C-Fus/TGA7–yEGFP and pBin19/Pro35S:TGA7–mGFP4 were transformed in yeast and agroinfiltrated in *N. benthamiana* leaves, respectively (Maier et al., 2011; Hermann et al., 2013). Fusion proteins were monitored by fluorescence microscopy and by immunodetection with an antiserum directed against GFP as described previously (Maier et al., 2011).

### RT-PCR

RNA isolation and RT-PCR analyses were done as reported by Zwicker et al. (2007). In each experiment shown as a separate panel in Figure 3 and in Supplementary Figure 9, RNAs were isolated from two or three individual plants or from two separate pools of seedlings. Altogether, results from six independent RNA samples isolated from young leaf tissue of six individual plants are shown in the figures. All experiments were repeated once with new RNA isolates and thus comprise at least four biological replicates for each RNA source. Reactions were run in parallel with and without reverse transcriptase (RT). The primer combinations and control plasmids used for the different gene fragments are listed in Table S1. When available, 1ng of plasmid DNA harboring the respective cDNA was subjected to PCR amplification in parallel with the RNA samples.

### Generation of transgenic tobacco plants, overexpression of *AtNPR1* in *N. benthamiana* and *GUS* reporter gene assays

Agrobacterium-mediated transformation of tobacco was performed as described (Grüner et al., 2003). Transgenic lines 378, carrying the *GUS* reporter gene under control of the upstream activating sequence of Gal4 transcription factor (*UAS^Gal4^: GUS*) and a gene for an artificial transcription factor (*Pro35S: GVG*; Figure S10), were generated by transformation with a vector developed by Aoyama and Chua (1997) and modified by Fan and Dong (2002). Regenerating plantlets were selected on hygromycin. Line 378-20, exhibiting intermediate GUS activity induced by DEX and no background activity in untreated and SA-treated leaf tissue (Figures S11A and S11B), was chosen for secondary transformation with pBin19/Pro35S: TGA7GBD. The resulting lines (380) were selected on medium containing hygromycin and kanamycin. Regenerated plants were tested for activity of the TGA7GBD chimeric protein (Figure S11C). Several independent primary transformants, all displaying SA-induced reporter gene expression, were selfed, and plants of the homozygous T3 generation were used for functional assays.

*Nicotiana benthamiana* plants were infiltrated with Agrobacteria harboring constructs pBin19/Pro35S:mGFP4 or pBin19/Pro35S:AtNPR1 as reported previously (Hermann et al., 2013). Proteins were extracted from agroinfiltrated leaf tissue 5 days post-infiltration when strong GFP fluorescence was observed.

Determination of GUS enzyme activity and histochemical localization of GUS activity in situ were performed as described by Weigel et al. (2001) and Glocova et al. (2005). GUS activities were measured in extracts directly sampled from tobacco leaf tissue or from leaf disks incubated on chemicals. Eight disks were collected from four leaf halves of young plants at the four- to six-leaf stage and floated on chemicals as indicated. Altogether, a minimum of three T3 plants of different lines 380 with the *TGA7GBD* chimeric gene, emerged from three independent transformation events, were tested for chemical induction. The experiments were repeated at least once with newly raised plants. Representative results obtained in one experimental set are shown. Protein extracts of whole seedlings were prepared from pools of ten individuals of the T3 generation deriving from three independent transformation events. The experiment shown in Supplementary Figure 11C was conducted at a later point of time to check previous results. In this case, proteins were extracted from three separate pools of seedlings for each line, and GUS activities are depicted as mean activities, plus and minus SD. Histochemical localization of GUS activity in T3 seedlings of lines 380 was performed with ten individuals each, emerged from four different transformation events. Representative results are shown. Protein extracts of older plant tissue were sampled from leaves of flowering plants at different positions on the axis (top, middle, bottom position) reflecting different developmental stages. Typically, leaves from the bottom part of the axis were yellowish, leaves from the middle third of the axis were fully grown and green, and leaves from the top of the axis close to the inflorescence were smaller and dark-green. Extracts were prepared from three plants each of three different transgenic lines 380. Transgenic line 138-3, containing a *Pro-1533PR-1a: GUS* construct (Grüner and Pfitzner, 1994), was used as control for SA-induced and senescence-associated *PR-1* gene expression. GUS activities are given in units (1 unit = 1nMol 4-MU formed per hour per mg protein).

### Generation of antisera against NPR1 proteins and immunodetection of proteins

*Arabidopsis NPR1* was excised as *Bam*HI/*Sal*I fragment from pGAD424 (Weigel et al., 2001) and ligated to expression vector pQE-32. *E. coli* overexpressed 6xHis–AtNPR1 purified over a Ni-NTA affinity resin was used for polyclonal antibody production in rabbits. Deletion *NtNPR1(386-588*), comprising the conserved C-terminal third of tobacco NPR1, was cloned as *Bam*HI fragment in pGEX-3X. *E. coli*-expressed GLUTATHIONE S-TRANSFERASE (GST)–NtNPR1(386-588) fusion protein was purified on a glutathione affinity column and used for immunization of rabbits.

For immunodetection of NPR1, proteins were extracted from tobacco leaves with GUS lysis buffer. Crude extracts were cleared by centrifugation, and equal amounts of protein or equal extract volumes were loaded on SDS polyacrylamide gels as indicated. Separated proteins were blotted, and nitrocellulose membranes were incubated with the a-6xHis–AtNPR1 serum. The same extracts used for immunodetection of NPR1 were also used for measuring GUS reporter enzyme activities. For detection of fusion proteins in yeast, extracts were prepared from transformed cells as reported by Weigel et al. (2001). Nitrocellulose filters were incubated with polyclonal antisera directed against *E. coli*-expressed Maltose-Binding Protein–NtNIMIN2a (MBP–NtN2a; Hermann et al., 2013), GST–NtNPR1(386-588) or 6xHis–AtNPR1. Protein loading was checked by staining nitrocellulose filters with Ponceau S (Hermann et al., 2013). Alternatively, unspecific bands reacting with the antisera are marked to demonstrate equal gel loading.

### Accession numbers

Sequence data from this article can be found in the EMBL/GenBank data libraries under accession numbers KY392756 (*NtTGA7*), XP_016476198 (*NtTGA1-like*), XP_009588814 (*NtoTGA1*), XP_006360289 (*StTGA7*), X16449 (*NtTGA1a*), M62855 (*NtPG13*), U90214 (*NtTGA2.1*), AF031487 (*NtTGA2.2*), AY998018 (*NtTGA10*), AT5G65210 (*AtTGA1*), AT5G06950 (*AtTGA2*), AT1G22070 (*AtTGA3*), AT5G10030 (*AtTGA4*), AT5G06960 (*AtTGA5*), AT3G12250 (*AtTGA6*), AT1G77920 (*AtTGA7*), AT1G68640 (*PAN*), AT1G08320 (*AtTGA9*), AT5G06839 (*AtTGA10*), XM_009763156 (*NsGRXC6*), AT4G33040 (*AtGRXC6*), AT1G28480 (*AtGRXC9*), AT1G64280 (*AtNPR1*), AT3G25882 (*AtNIMIN2*), AF480488 (*NtNPR1*), AY640382 (*NtNIM1-like1*), AF057379 (*NtNIMIN2a*), EF015598 (*NtNIMIN2c*), and X06930 (*NtPR-1a*).

## RESULTS

### Structure of tobacco TGA7

In a Y2H screen with the AtNPR1 bait, we isolated three clones derived from the same gene coding for a TGA transcription factor. The assembled cDNA sequence of this gene encodes a 363 amino acid-long novel member of the tobacco TGA family. The domain structure of the protein, as compared to the structures of two other family members, NtTGA1a and NtTGA10, is shown in Figure S1. The bZIP region (41 amino acids) of the novel TGA factor is preceded by an 80 amino acid variable N-terminus and followed by a conserved C-terminal region. The conserved C-terminus encompasses 242 amino acids, including two glutamine-rich stretches, Q1 and Q2 (Katagiri et al., 1989), which are common to all family members from tobacco and *Arabidopsis*. The amino acid sequence is identical to predicted proteins TGA1-like from *N. tabacum* and *N. tomentosiformis* (Nto), and 91% identical to predicted protein TGA7 from *Solanum tuberosum* (St). Among known *N. tabacum* TGA factors, the novel factor is most similar to TGA1a exhibiting 53% identity (Figures S2 and S3A). We also compared the sequence of the novel tobacco protein to *Arabidopsis* TGA factors, because, in *Arabidopsis*, the whole family is known. It stands between *Arabidopsis* clade I (AtTGA1, AtTGA4) and clade III (AtTGA3, AtTGA7) factors (Figure S3B and S3C). Based on its close relatedness to StTGA7 and to *Arabidopsis* clade III factors, and its obvious disparity to NtTGA1a, the newly identified factor was named NtTGA7. The GenBank accession number of TGA7 is KY392756.

### Transactivation activity of TGA7 in yeast

To test the DNA binding potential of TGA7, we used a yeast one-hybrid (Y1H) system (Wolf et al., 1996). One copy of the *as-1* cis-acting element from the *CaMV 35S* promoter was cloned in a vector upstream of the *CYC1* minimal promoter fused to the *lacZ* gene. The construct was transformed in yeast cells together with vectors containing fusion genes of various TGA factors with the Gal4 transcription activating domain (GAD). In this system, binding of GAD fusion proteins to the *as-1* DNA element produces β-galactosidase reporter activity.

We observed rather low level reporter gene activity already in absence of TGA factors (**Figure 1**). This is likely due to binding of endogenous yeast bZIP proteins, e.g., GCN4 activator, to the *as-1* sequence. Reporter enzyme activity was, however, significantly increased upon expression of *Arabidopsis TGA2* and *TGA6*, and tobacco *TGA2.1*, *TGA2.2* and *TGA10* fusion genes. In comparison to these factors, GAD–TGA7 produced only very low reporter gene activity, just slightly above the background. Inability to mediate substantial β-galactosidase enzyme activity is, however, not due to lack of accumulation of the fusion protein since GAD-TGA7 yielded weak activity above the background level in all Y1H assays and strong reporter activity in diverse Y2H assays performed by us (see, f.e., Figure S6). Among the tobacco TGA factors tested, transactivation of TGA7 is only marginal in an *as-1*-based Y1H system. Our finding may reflect lack of strong DNA binding activity of TGA7 to *as-1*. Alternatively, endogenous bZIP proteins may obscure interaction of GAD–TGA factors with DNA in yeast.

**Figure 1.**
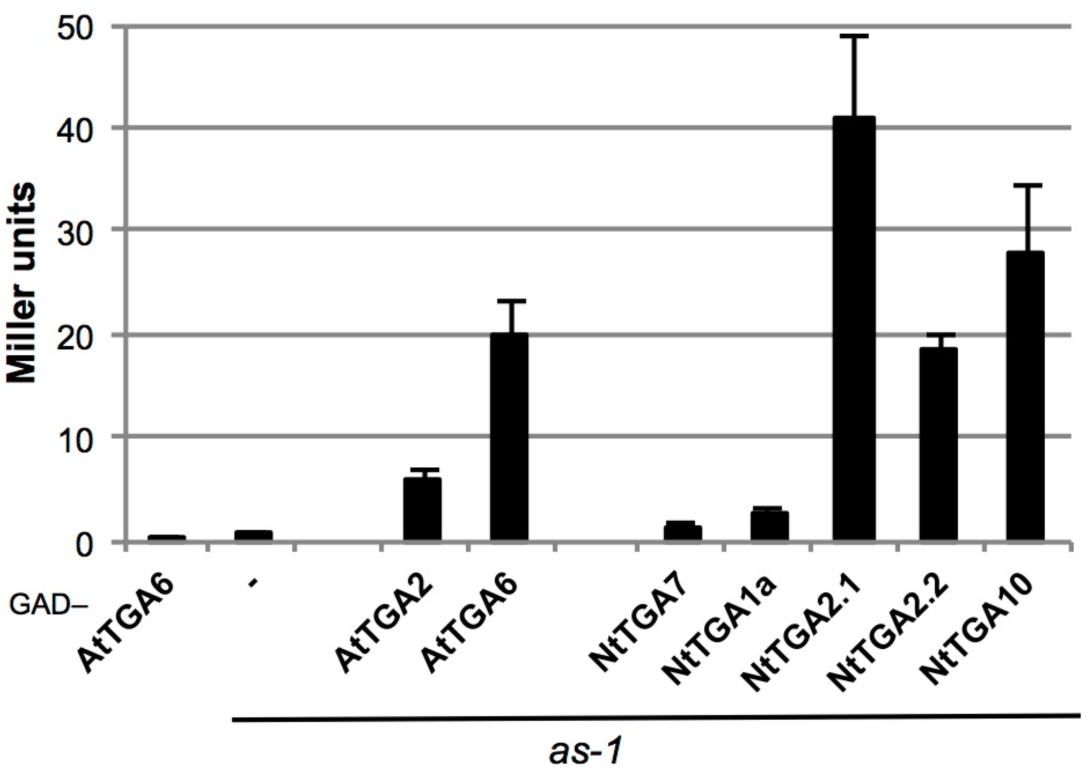
Test for binding of TGA7 to the *as-1* cis-acting element. Binding of TGA7 to one copy of *as-1* from the *CaMV 35S* promoter was determined in quantitative Y1H assays. *TGA* factor genes were expressed as fusions with the sequence for GAD. Activity of TGA7 is compared to the activities of *Arabidopsis* TGA2 and TGA6, and to the capacities of tobacco TGA1a, TGA2.1, TGA2.2 and TGA10. The experiment was repeated once. Representative results are shown.

In our previous work, we have shown already that TGA7 (termed NtTGA8 in Maier et al., 2011) exhibits transcriptional activity in Y1H assays when fused to the Gal4 DNA-binding domain (GBD), and that addition of SA to yeast growth medium does not affect this activity of TGA7. The level of transcriptional activity of TGA7 in yeast is similar to that of its close relative NtTGA1a (Maier et al., 2011; Figure S4B). Furthermore, as reported for other TGA transcription factors, e.g., NtTGA1a and NtTGA2.1 (Neuhaus et al., 1994; Pascuzzi et al., 1998; Niggeweg et al., 2000a), transcriptional activity is mediated by the variable N-terminus of TGA7. Deletion mutant GBD-TGA7Δ1, lacking the whole N-terminus including the basic DNA-binding domain, mediates only residual transcription activity in yeast (Figures S4 and S5).

TGA factors are known to form dimers among TGA family members and with glutaredoxins. Consistent data were obtained with TGA7. TGA7 is able to form homodimers as well as heterodimers with other TGA factors. Homodimerization of TGA7 and heterodimerization with NtTGA1a or NtTGA2.1 appear to be preferred over binding to NtTGA2.2 in yeast (**Figure 2A**; Figure S6). Similar results were obtained with the close relative NtTGA1a (**Figure 2B**). In contrast, NtTGA2.2 clearly shows another profile by preferential interaction with NtTGA2.1 (**Figure 2C**). We also tested the impact of the leucine zipper motif on dimer formation. It is commonly assumed that heptad leucine zippers create an amphiphatic helix, and that TGA factors interact with each other via the hydrophobic sides of these helices. Surprisingly, dimerization of TGA7 does not require two zipper motifs. Deletion mutant TGA7Δ2, lacking the leucine zipper (Figure S4A), supports homodimerization with TGA7Δ1 as well as heterodimerization with NtTGA1aΔ1 to similar extents as determined for homodimers of TGA7Δ1 and NtTGA1aΔ1, respectively, which possess intact zipper motifs (Figures S5A and S7). Similarly, Boyle and associates showed that AtTGA2 lacking 93 amino acids from the N-terminus including the bZIP region is still able to homodimerize (Boyle et al., 2009). Together, the data establish that leucine zipper motifs are not needed for association of TGA factors. Instead, a dimer stabilization (DS) region identified in the conserved C-terminus of NtTGA1a may be sufficient to form homo- and heterodimers among TGA transcription factors (Katagiri et al., 1992).

**Figure 2.**
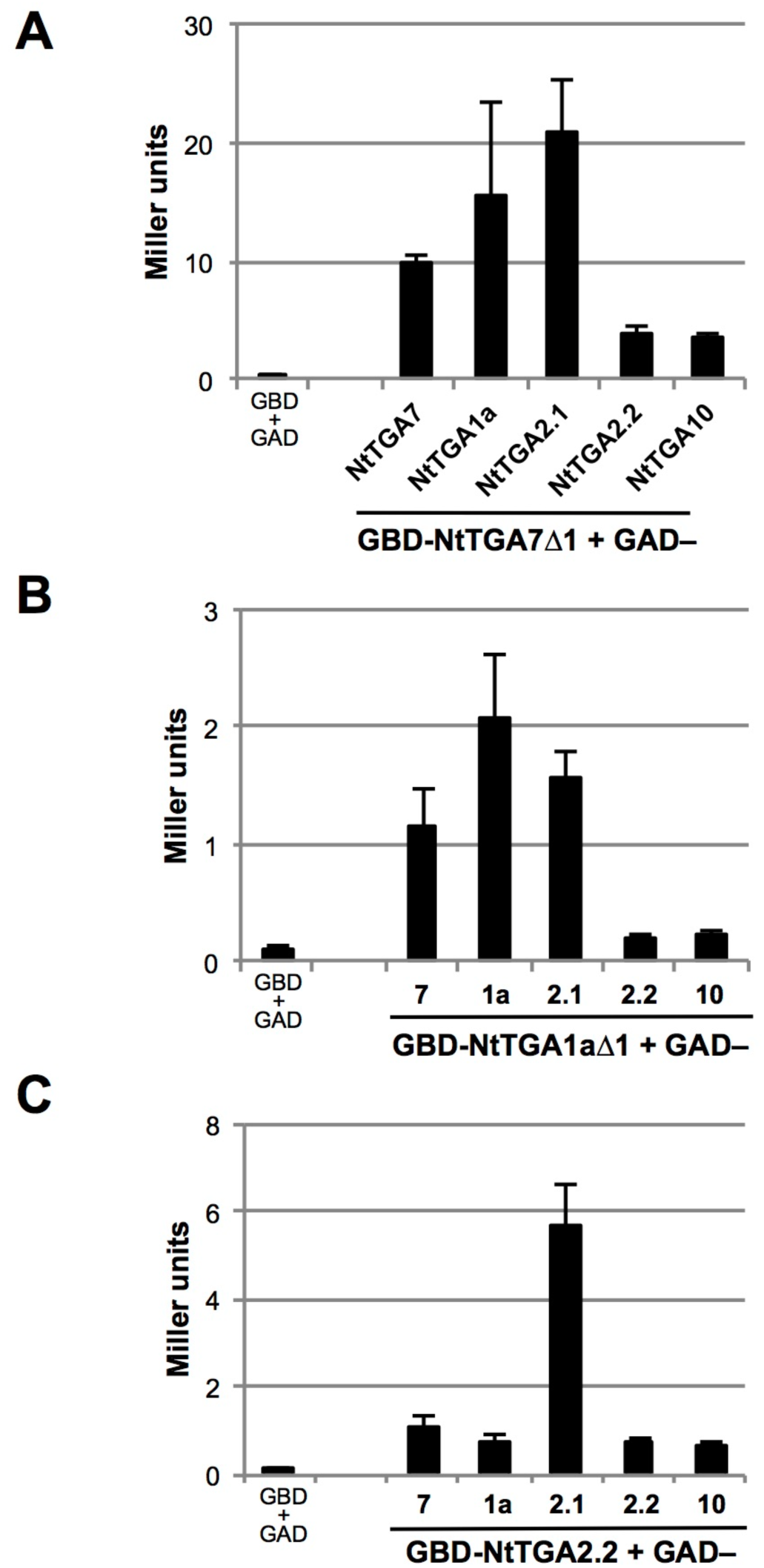
TGA7 interacts differentially with TGA factors. **(A)** Dimerization of TGA7Δ1 with tobacco TGA factors. Interaction of TGA7Δ1 with NtTGA family members was determined in quantitative Y2H assays and compared to the dimerization potentials of NtTGA1aΔ1 (**B**) and NtTGA2.2 (**C**). For transcription activators TGA7 and NtTGA1a, truncated versions Δ*1*, lacking own transcription activity, were expressed as fusions with the sequence for GBD. Factor NtTGA2.2 does not exhibit transcription activity in yeast. The experiment was repeated once. Representative results are shown.

In yeast, TGA7 also interacts with glutaredoxins from tobacco and *Arabidopsis*. In a Y2H screen using TGA7Δ1 as bait, we identified GLUTAREDOXIN (GRX)-C6 coding for a tobacco CC-type glutaredoxin, which is identical to GRXC6 predicted from the *N. sylvestris* (Ns) genome (V.S. and U.M.P., unpublished). The *Arabidopsis* ortholog AtGRXC6, synonym ROXY21, interacts with AtTGA2 in Y2H assays (Zander et al., 2012). Likewise, AtGRXC6 and AtGRXC9 (synonyms ROXY19/GRX480) interact with TGA7 (Figure S5B). Furthermore, deletion mutant TGA7Δ2, without leucine zipper motif, is sufficient for interaction with NtGRXC6 (Figures S5A and S7).

### Subcellular localization of TGA7–GFP fusion proteins

Consistent with their functional role as transcription activators, TGA factor fusion proteins have been shown to accumulate in nuclei in transgenic tobacco plants (Van der Krol and Chua, 1991). Accordingly, we detected TGA7–GFP fusion proteins in nuclei of yeast cells and in nuclei of *N. benthamiana* epidermal cells (Figure S8).

### Expression of *TGA7*

*TGA7* mRNA levels were analyzed in different organs from tobacco plants by reverse transcription-polymerase chain reactions (RT-PCR). Expression of *TGA7* was compared to expression of TGA factors *NtTGA2.2* and *NtTGA1a* as internal controls. *NtTGA2.2* is expressed moderately in leaves and in roots (Niggeweg et al., 2000a), while *NtTGA1a* is predominantly expressed in roots (Katagiri et al., 1989). *TGA7* expression was also compared to expression of the pathogen defense marker *PR-1a*. RNAs were isolated from different parts of tobacco plants or after different treatments of leaf tissue, and equal amounts of RNAs were subjected to RT-PCR. The primer pairs used and the sizes of fragments generated by PCR from plasmids carrying cDNAs for *TGA7* and several control genes are listed in Table S1. Surprisingly, we were not able to detect *TGA7* mRNA in young leaf tissue expressing *NtTGA2.2* (**Figure 3A**; Figure S9). Yet, *TGA7* transcripts were evident in whole seedlings (**Figure 3C**). The gene is neither induced by treatment of young leaves with SA nor in the course of the pathogen defense response HR elicited by infection with *Tobacco mosaic virus* (*TMV*; Figure S9). However, *TGA7* mRNA was present in tissue from older leaves of flowering plants (**Figure 3A**). In young plants, *TGA7* transcripts were readily found in petiole, stem and root containing high contents of vascular tissue (**Figure 3**).

**Figure 3.**
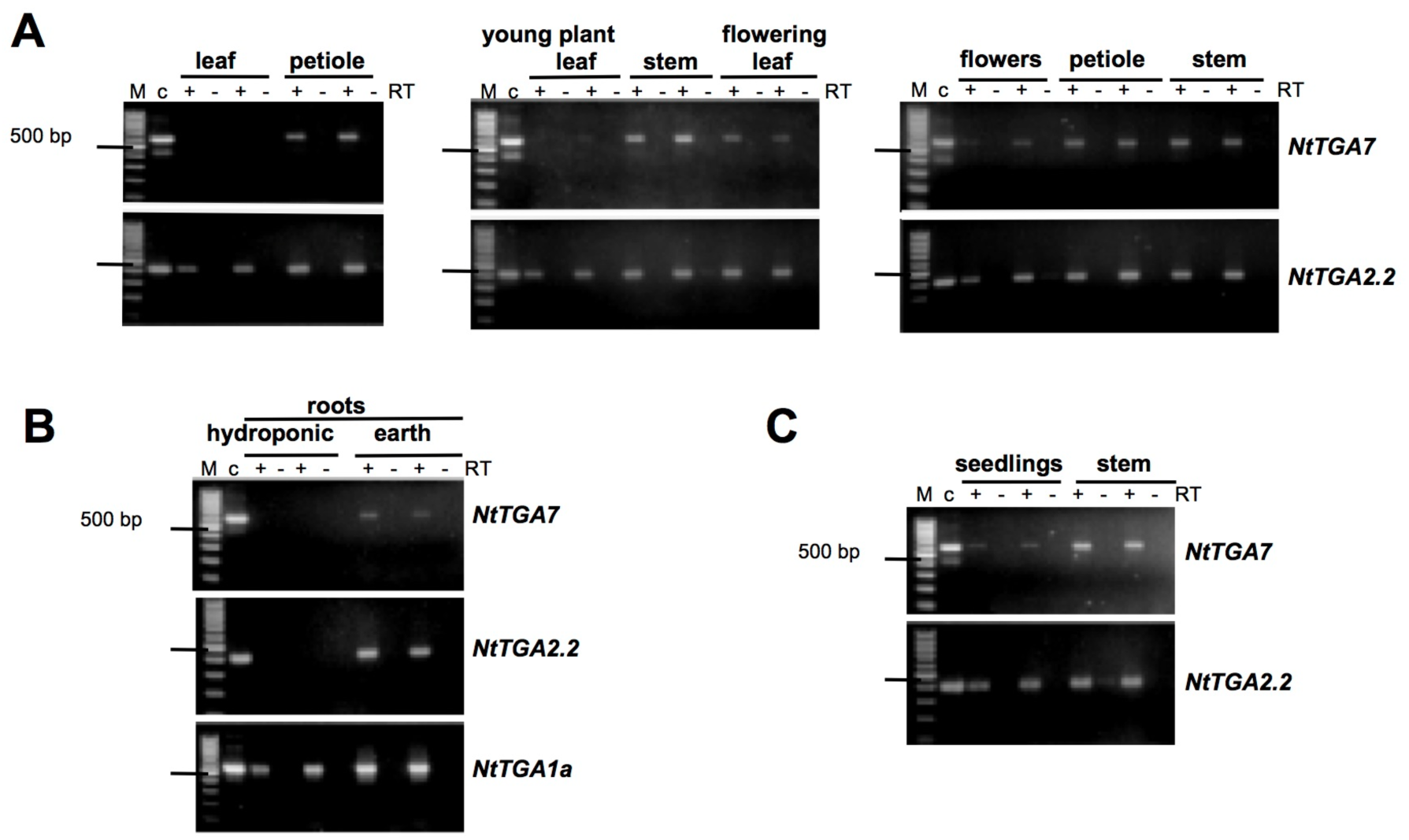
Expression of *TGA7*. RNA samples from different organs of tobacco plants and from whole seedlings were analyzed by RT-PCR. RNAs were isolated from two individuals or two individual pools in each experiment shown as a separate panel. Expression of *TGA7* is compared to expression of tobacco family members *TGA2.2* and *TGA1a*. RT-PCR analyses were performed in absence and presence of RT with primer combinations listed in Table S1. In lanes c, PCR products from 1ng of plasmid DNAs carrying the respective cDNAs were loaded. The experiments were repeated with RNAs isolated from different sources. Representative results are shown. **(A)** Expression of *TGA7* in aboveground organs of young and flowering plants. **(B)** Expression of *TGA7* in roots of young plants. **(C)** Expression of *TGA7* in whole seedlings.

### TGA7 acts as salicylic acid-dependent transcriptional acivator in planta

Since TGA7 is able to activate transcription in Y1H assays, we asked whether TGA7 would also be a transactivator in tobacco plants. We used a chimeric reporter gene system developed by Aoyama and Chua (1997), which had shown that AtTGA2 serves as an SA-activated and NPR1-dependent transcription factor in *Arabidopsis* (Fan and Dong, 2002). The system is depicted in Figure S10. *TGA7* was modified as described by Fan and Dong (2002). The sequence coding for the bZIP region of TGA7 was replaced by the sequence coding for Gal4 DNA-binding domain. The resulting construct, encoding a TGA7GBD chimera, was put under control of the *CaMV 35S* promoter and stably integrated into an *N. tabacum* reporter line containing a genomic construct with the structural gene for β-glucuronidase (*GUS*) fused to a synthetic promoter consisting of a minimal *CaMV 35S* promoter sequence and six copies of the Gal4 DNA-binding site (*UAS^GAL4^: GUS*). The transgenic reporter line also contained the gene for an artificial transcription factor, termed GVG, driven by the *CaMV 35S* promoter (Pro*35S: GVG*). GVG comprises the GBD (G), the VP16 transactivation domain (V), and a glucocorticoid binding site (G). In presence of steroid hormones, e.g., dexamethasone (DEX), cytoplasmic GVG is translocated to the nucleus where it binds to *UAS^GAL4^* and activates transcription of the *GUS* reporter. In transgenic lines with the GUS reporter and the *GVG* gene (lines 378), we detected only marginal GUS activity in absence of DEX and in presence of SA, but high levels of enzyme activity after exposure of plant tissue to DEX (Figures S11A and S11B). Hence, GUS activity in absence of steroids would be due to reporter gene transcription mediated by the TGA7GBD fusion protein in plant lines 380.

Leaf tissue of ten independent primary transformants (T0 generation; lines 380) containing the *TGA7GBD* construct in the *GVG+ UAS^Gal4^:GUS* genomic context of line 378-20 was analyzed. In all cases, we determined very high levels of GUS reporter enzyme activity in presence of DEX in lines 380 (Figure S11C). Seven transformants exhibited low levels of background GUS activity without DEX. This reporter activity was, however, significantly increased upon treatment of plant tissue with SA showing that the chimeric factor TGA7GBD activates transcription in tobacco leaves in response to the SAR signal SA (average GUS induction by SA 14.5-fold).

Primary transformants with the *Pro35S: TGA7GBD* construct were selfed, and further analyses were performed with T3 plants from a total of six independent transgenic lines. Chemical induction was conducted with leaf tissue from young tobacco plants at the four- to six-leaf stage. All individuals tested exhibited responsiveness to SA (11.9-fold average GUS induction by SA; ten biological replicates; **Figures 4A** and **4B**; Figure S12A). Transcription of *GUS* was likewise increased with the SA analog benzo(1,2,3)thiadiazole-7-carbothioic acid S-methyl ester (BTH), which induces *PR-1* genes and SAR in wheat, *Arabidopsis* and tobacco (Görlach et al., 1996; Friedrich et al., 1996; 16.7-fold average GUS induction by BTH; seven biological replicates; **Figures 4A** and **4B**). The non-functional SA analog 4-hydroxy benzoic acid (4-OH BA; White, 1979) could not activate TGA7GBD (three biological replicates; **Figure 4B**). Similarly, other phytohormones, like indole-3-acetic acid (IAA), 1-naphthalene acetic acid (NAA), 2,4-dichlorophenoxy acetic acid (2,4-D), gibberellic acid (GA), and methyl jasmonate (MeJA), had no effects on TGA7GBD (Figure S12A and data not shown). MeJA is an inducer of *PR* genes other than *PR-1* and acts as an antagonist of SA-induced defense and *PR-1* gene expression (Spoel et al., 2003; Pieterse et al., 2012). We therefore tested the effects of simultaneous application of both SA and MeJA. In fact, activation of the chimeric transcription factor was not observed in presence of SA and MeJA in leaf tissue (three biological replicates; Figure S12A). These data further support the view that TGA7 serves as part of a transcription complex in tobacco that is positively linked to SA signaling. Of note, TGA7GBD mediates only very weak reporter gene activity in untreated leaf tissue from young plants. However, we found considerable GUS reporter enzyme activity in whole seedlings germinated on MS medium with or without addition of sucrose (six biological replicates; **Figure 4C**; Figures S12B and S12C). This actitvity is strictly localized to cotyledons and primary leaves of seedlings (**Figure 4D**), and thus is due to activity of TGA7GBD, but not to activity of the GVG transcription factor, which produces reporter activity in both leaves and root tips only in presence of DEX (Figure S11B). Importantly, seedling-specific transactivation of the reporter by TGA7GBD was not paralleled by expression of the SAR marker *PR-1* (Figure S12D; **Figure 4C**). Similarly, we observed considerable GUS reporter enzyme activity in leaf tissue from older plants carrying inflorescences without prior application of SA or pathogen infection (nine biological replicates; **Figure 4E**). This particular pattern of TGA7GBD activity is reminiscent of induction of *PR-1a* gene transcription in mature tobacco plants (**Figure 4E**; Figure S12E). *PR-1a* is activated in leaf tissue from flowering plants, while it is inactive in leaves from young plants at the four- to six-leaf stage, where *PR-1a* is strongly induced in the course of HR and in response to SA (Grüner and Pfitzner, 1994). Indeed, GUS reporter activities in old leaf tissue from *Pro35S: TGA7GBD* transgenic plants were similar to GUS activities in plants with the *ProPR1a: GUS* construct (**Figure 4E**). Interestingly, *TGA7* transcripts were clearly detected in leaves of flowering plants and in whole seedlings, although, as compared to the expression of *NtTGA2.2*, they appeared not as abundant as in stems of young plants (**Figures 3A** and **3C**).

**Figure 4.**
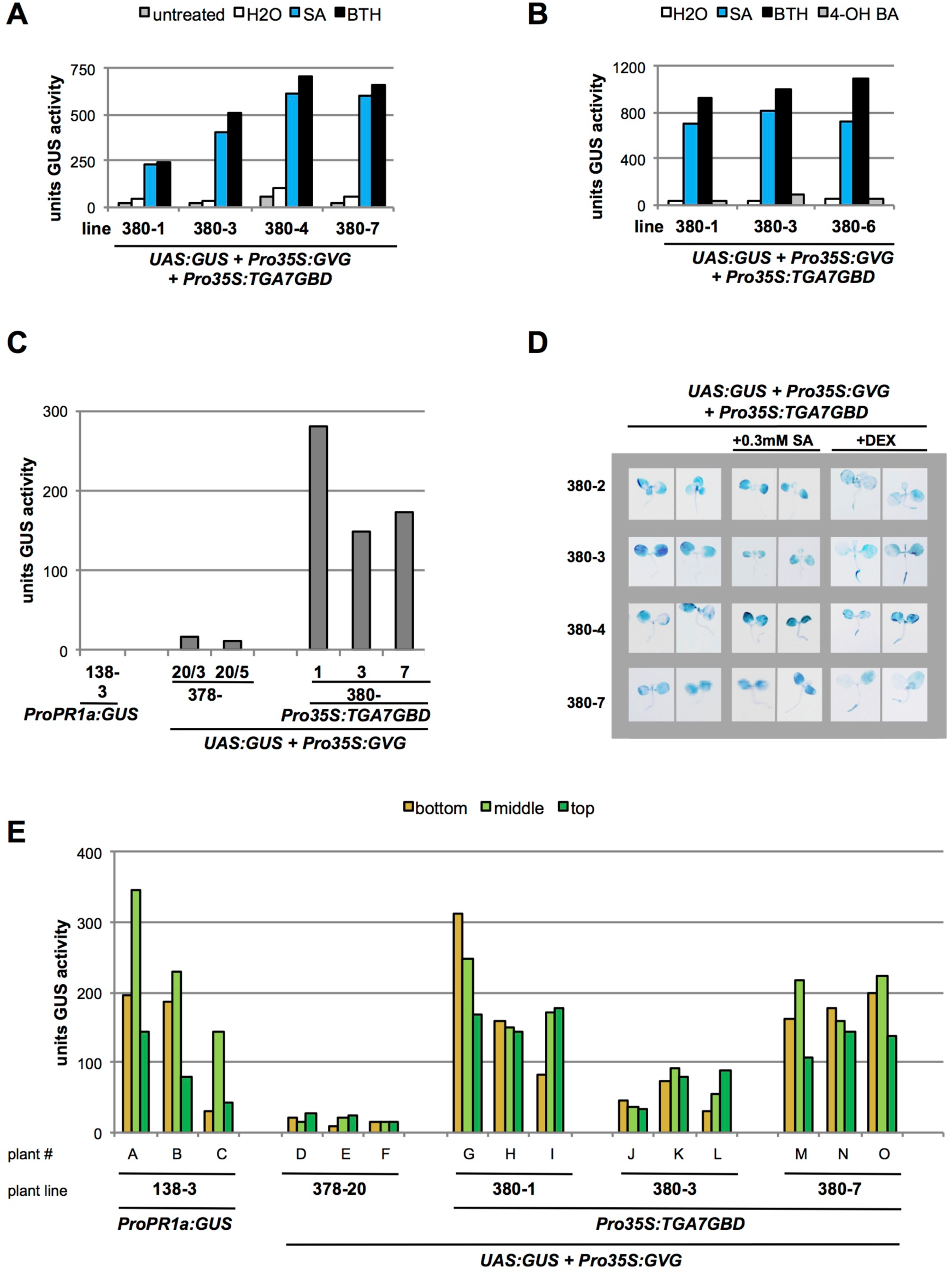
A TGA7GBD chimeric protein mediates SA-dependent and SA-independent gene activation in tobacco. Transgenic line 378-20, which harbors a reporter construct with *GUS* under control of the upstream activating sequence of Gal4 transcription factor (*UAS^Gal4^: GUS*) and an effector construct with an artificial transcription activator gene including the sequence for the Gal4 DNA-binding domain (*Pro35S: GVG*), was transformed with a vector containing the *Pro35S: TGA7GBD* chimeric gene yielding lines 380. **(A)** GUS enzyme activity in salicylic acid and BTH-treated young leaves. GUS activity was determined in four independent transgenic lines (380-1, 380-3, 380-4 and 380-7). The experiment was repeated once. Representative results are shown. **(B)** GUS enzyme activity in young leaves treated with salicylic acid, BTH or 4-hydroxybenzoic acid. GUS activity was determined in three independent transgenic lines. The experiment was repeated once. Representative results are shown. **(C)** GUS enzyme activity in four-week-old seedlings grown on MS medium. GUS activities in three independent transgenic lines with the *TGA7GBD* construct are compared to GUS activities in the parental line 378-20 with the GVG transactivator and to GUS activities in a line carrying the *Pro-1533PR1a:GUS* construct. GUS activities were determined from pools of ten seedlings for each measurement. **(D)** Histochemical localization of GUS enzyme activity in three to four-week-old seedlings grown on MS medium without addition or supplemented with salicylic acid or dexamethasone. GUS activity was determined in four independent transgenic lines. **(E)** GUS enzyme activity in leaves of different developmental stages. GUS activities in three plants each of three independent transgenic lines with the *TGA7GBD* construct are compared to GUS activities in the parental line 378-20 with the GVG transactivator and to GUS activities in a line carrying the *Pro-1533PR1a: GUS* construct. Leaf tissue from bottom, middle or top positions on the axes of older tobacco plants carrying inflorescences was analyzed.

### TGA7 interacts preferentially with the NtNPR1 C-terminal end

We identified *TGA7* in a Y2H screen using AtNPR1 as bait. In total, three independent cDNA clones were isolated in this screen, indicating that binding of TGA7 to AtNPR1 must be specific and significant. Surprisingly, the amino acid sequence of *TGA7* revealed two cysteines, Cys-255 and Cys-261, in the same positions as found in AtTGA1 and AtTGA4 (Figure S13). The cysteine residues have been reported to preclude binding of Arabidopsis TGA1 and TGA4 to AtNPR1 in Y2H assays (Després et al., 2003). This apparent discrepancy prompted us to analyze interaction between TGA7 and NPR1 in more detail.

As opposed to NtTGA1a and NtTGA10, TGA7 indeed binds to AtNPR1 in Y2H assays, although interaction with TGA7 appears weaker than interactions with NtTGA2.1, NtTGA2.2 or AtTGA2 (Figure S14). In contrast to the AtNPR1 interaction profile, however, none of the tobacco TGA factors exhibited clear binding to GBD–NtNPR1 in qualitative Y2H *His* reporter gene assays (data not shown). In quantitative β-galactosidase activity assays, we detected low level interaction of GBD–NtNPR1 with GAD–NtTGA2.1, GAD–NtTGA2.2 and GAD–TGA7, respectively. With these factors, *lacZ* activities were higher than transcriptional activity of GBD–NtNPR1 alone, which we had already observed in our previous work (**Figure 5A**; Maier et al., 2011).

**Figure 5.**
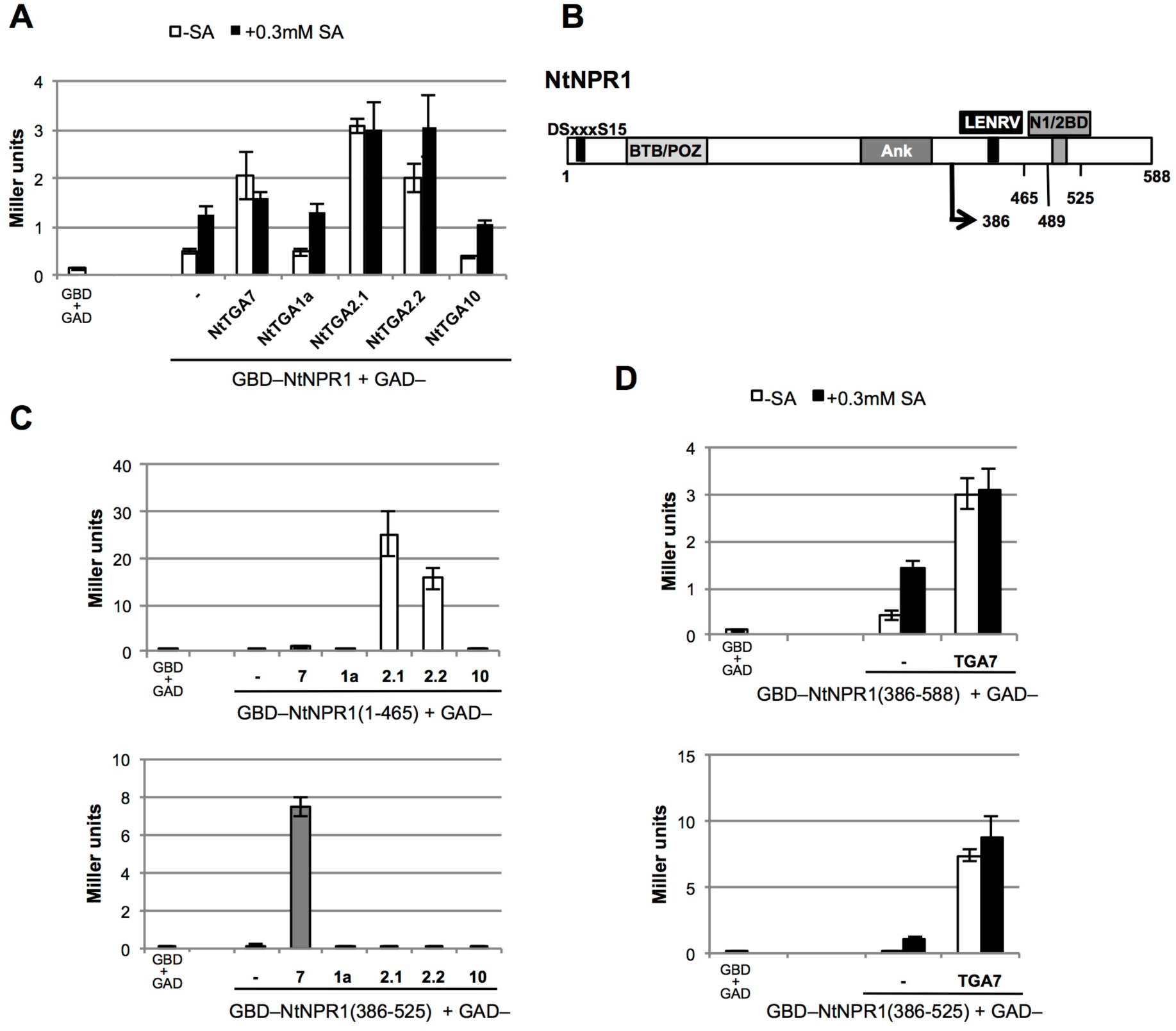
TGA7 interacts with the C-terminus of tobacco NPR1. Interaction of TGA7 with NtNPR1 full-length and truncated proteins, as determined in quantitative Y2H assays, is compared to interactions with other tobacco TGA family members. The experiments were repeated once (A) or twice (C,D). Representative results are shown. **(A)** Interaction of tobacco TGA factors with NtNPR1 full-length protein in absence and presence of SA. **(B)** Domain structure of tobacco NPR1. The LENRV motif and the binding domain for NIMIN1 and NIMIN2-type proteins (N1/2BD), involved in SA sensing by NtNPR1, are indicated. Other conserved domains are a phosphodegron motif at the N-terminus, a BTB/POZ domain and an ankyrin repeat region. Positions of deletion mutants are marked. **(C)** Interaction of tobacco TGA factors with NtNPR1 deletions encompassing amino acids 1 to 465 and amino acids 386 to 525, respectively. **(D)** Interaction of TGA7 with truncated NtNPR1 proteins comprising C-terminal regions 386 to 588 or 386 to 525 in absence and presence of SA.

Attempting to map the TGA interaction site on NtNPR1, we used several deletion constructs. Three constructs were informative (**Figure 5B**). Both clade II factors, NtTGA2.1 and NtTGA2.2, bind strongly to truncated protein NtNPR1(1-465), lacking the C-terminus (**Figure 5C**). This finding is consistent with previous observations suggesting that *Arabidopsis* NPR1 binds TGA factors predominantly via its central ankyrin repeat region. Quite unexpectedly, TGA7 binds differently to NtNPR1. Interaction of TGA7 with NtNPR1(1-465) is marginal when compared to interaction levels of NtTGA2.1 and NtTGA2.2. Conversely, TGA7 is able to bind to the C-terminal end of NtNPR1 from positions 386 to 525 (**Figure 5C**). Co-expression of the other *TGA* factor genes produces no *lacZ* activity with NtNPR1(386-525). TGA7 is also able to bind to truncated protein NtNPR1(386-588), but not to NtNPR1(386-489) (**Figure 5D**; Figures S15A and S15B). Thus, among five tobacco TGA factors tested, TGA7 is exclusive in that it interacts with the C-terminal region of NtNPR1. Yet, although TGA7 binds to the respective full-length proteins (Figure S14; Maier et al., 2011), it does not bind to the NtNIM1-like1 or AtNPR1 C-termini (Figure S16 and data not shown). Our finding is noteworthy, since the C-termini of NPR1 proteins from tobacco and *Arabidopsis* have been shown to harbor motifs for sensitivity of NPR1 toward the SAR signal SA and also for binding SA-induced NIMIN proteins (Maier et al., 2011). Binding of TGA7 to the tobacco NPR1 C-terminus does not, however, seem to be affected by SA (**Figure 5D**). Likewise, interaction of other TGA factors with NtNPR1 is not influenced significantly by SA (**Figure 5A**; Maier et al., 2011). In this respect, TGA7 clearly differs from binding of NtNIMIN2-type proteins to the NtNPR1 C-terminus which is impaired in presence of SA (Maier et al., 2011).

We also undertook efforts to narrow down the binding region for NtNPR1 on TGA7. TGA7Δ2 (Figure S4A) is still able to interact significantly with NtNPR1 deletions 386-525 and 386-588 (Figures S15C and S15D). Hence, the conserved C-terminal region of TGA7 binds different partner proteins, TGA factors, glutaredoxins, and NPR1, while the variable N-terminus mediates transcriptional activity of TGA7, and the basic region may allow sequence specific contacts with DNA.

### Presence of NIMIN2-type proteins compromises binding of TGA7 to the NtNPR1 C-terminus

NIMIN-type proteins have been identified through Y2H screens with different NPR1 bait proteins (Weigel et al., 2001; Chern et al., 2005; Zwicker et al., 2007). Recent evidence indicates that they act as regulators of NPR1 activity and SAR in *Arabidopsis*, rice and tobacco (Weigel et al., 2005; Chern et al., 2005; Zwicker et al., 2007; Hermann et al., 2013). Binding of NIMIN2 has been located to a highly conserved domain, extending from positions 494 to 510 in the NtNPR1 C-terminus (**Figure 5B**; Maier et al., 2011), and thus might interfere with TGA7 interaction which occurs between positions 386 and 525. We therefore analyzed interactions of TGA7 and NIMIN2 proteins with the NtNPR1 C-terminus.

Initially, we tested whether NtNPR1 can interact simultaneously with both NIMIN2 and TGA7 in yeast cells. Previously, we have demonstrated that ternary protein complexes can assemble on *Arabidopsis* NPR1 with AtNIMIN1, AtNIMIN2 or AtNIMIN3 and AtTGA2 or AtTGA6 (Weigel et al., 2001). TGA7 was expressed as fusion with GAD, and NIMIN2-type proteins, NtNIMIN2a and NtNIMIN2c, were expressed as GBD fusions. TGA factors are not able to interact with NIMIN proteins (Weigel et al., 2001; Chern et al., 2014). NPR1 was put under control of the *MET25* promoter which is de-repressed in absence of methionine in yeast growth medium (Tirode et al., 1997). We were not able to detect complex formation between NtNPR1, TGA7 and NtNIMIN2 proteins in yeast (data not shown). This failure could be due to competition of TGA7 and NIMIN2 for the NtNPR1 C-terminus. Indeed, constitutitve interaction between GBD–NtNPR1 or GBD–NtNPR1(386-588) and GAD–TGA7 was disrupted in presence of NtNIMIN2a or NtNIMIN2c expressed from the *MET25* promoter (**Figure 6A**). High levels of NtNIMIN2a (**Figure 6B**) completely abolished binding of TGA7 to NtNPR1, while addition of SA to yeast culture medium partially restored the NtNPR1– TGA7 interaction due to SA sensitivity of NIMIN2 binding to the NtNPR1 C-terminus (**Figure 6A**; Maier et al., 2011). Thus, NtNPR1–TGA7 interaction occurs with and without SA, but presence of NIMIN2 compromises TGA7 binding to NtNPR1. Unfortunately, we were not able to map the exact binding site for TGA7 on the NtNPR1 C-terminus. Yet, TGA7 interaction is not abolished in the NIMIN2 binding mutant NtNPR1 F505/506S (Maier et al., 2011). Therefore, TGA7 does not appear to bind to the very same site on NtNPR1 as NIMIN2-type proteins. We do not exclude, however, that TGA7 binds to NtNPR1 in close proximity to the NIMIN2 interaction site.

**Figure 6.**
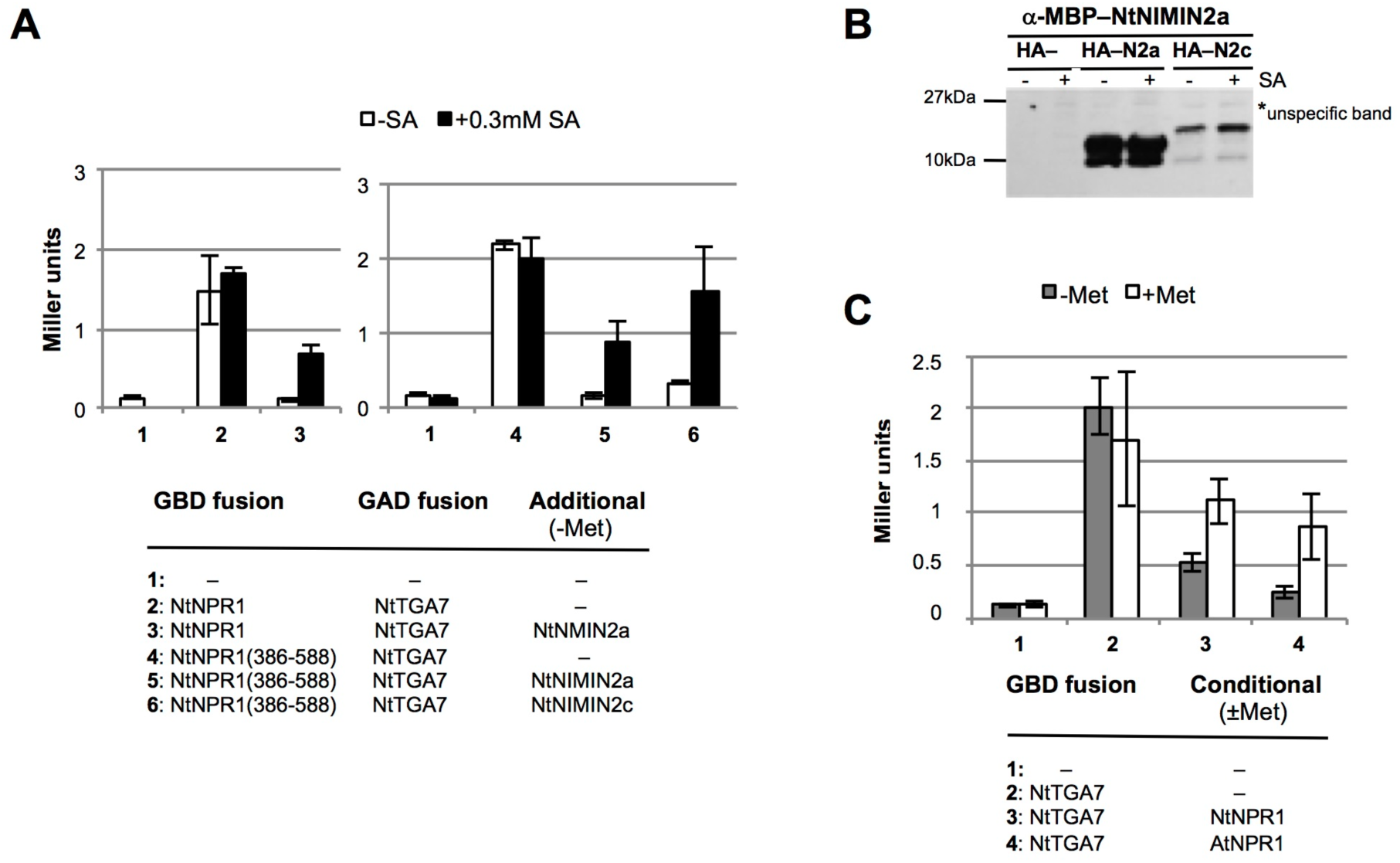
Binding of TGA7 to tobacco NPR1 is compromised by NIMIN2 proteins, and TGA7 transcription activity is compromised by interaction with NPR1. The experiments were repeated twice (A) or once (C). Representative results are shown. **(A)** Interaction of TGA7 with NtNPR1 in presence of NtNIMIN2-type proteins. Interactions were monitored in Y3H assays without and with SA. *NIMIN2* genes were expressed from the *MET25* promoter in absence of methionine. **(B)** Accumulation of NtNIMIN2-type proteins in yeast. NtNIMIN2a and NtNIMIN2c fusion proteins were detected with an antiserum raised against Maltose-Binding Protein (MBP)–NtNIMIN2a. **(C)** Transcription activity of TGA7 in presence of NPR1. Activity of TGA7 was monitored in yeast. Tobacco and *Arabidopsis NPR1* were expressed from the *MET25* promoter which is de-repressed in absence and repressed in presence of methionine.

### Interaction with NPR1 Masks Transcription Activity of TGA7

We have shown that NtNPR1 binds transcriptional activator TGA7 via its C-terminus, and that this interaction is outcompeted by presence of NIMIN2 proteins. Furthermore, we have demonstrated that transcription factor TGA7 is able to activate gene expression in tobacco leaves in response to SA. Together, these data may suggest that TGA7 transcription activity could be unleashed in planta by interplay of NtNPR1 with SA-induced NIMIN2 proteins.

To test whether TGA7 activity can be affected directly by interaction with NPR1, we expressed *TGA7* as fusion with GBD in yeast cells. For interaction studies, we used tobacco and *Arabidopsis* NPR1 since TGA7 binds differently to the two proteins. Co-expression of both *NtNPR1* and *AtNPR1* from the *MET25* promoter clearly dampened *lacZ* expression irrespective of their mode of interaction with TGA7. On the other side, inhibition of TGA7 transcription activity was partially relieved upon repression of *NPR1* by addition of methionine to yeast culture medium (**Figure 6C**).

### NPR1 protein accumulates constitutively in tobacco leaf tissue

We also monitored occurrence of NPR1 protein in tobacco plants at different physiological states. Recent reports have described that NPR1 is targeted by multiple post-translational modifications in *Arabidopsis* that control NPR1 accumulation and induction of defense genes (Mou et al., 2003; Tada et al., 2008; Lee et al., 2015; Saleh et al., 2015; Withers and Dong, 2016). Thus, NPR1 levels were found to rise significantly after treatment of *Arabidopsis* plants with SA (Fu et al., 2012). While this finding is consistent with the positive role *NPR1* plays in the SAR response based on genetic data (Cao et al., 1997; Ryals et al., 1997; Shah et al., 1997; Canet et al., 2010), other reports have suggested that there is no strict correlation between amounts of NPR1 protein on the one hand and levels of pathogen resistance on the other hand in *Arabidopsis* (Cao et al., 1998; Friedrich et al., 2001).

We analyzed NPR1 accumulation in crude protein extracts from tobacco using a polyclonal antiserum raised against *Arabidopsis* 6xHis–NPR1. In extracts from *Arabidopsis* leaves, the a-6xHis–AtNPR1 serum detects a major band co-migrating with the 70 kDa marker protein consistent with the calculated molecular weight of AtNPR1 of 66 kDa. In addition, the serum yields a faint band slightly above the 70 kDa marker. Essentially the same pattern was observed with leaf extracts from *N. tabacum* and *N. benthamiana* (**Figure 7A**; Figure S17A). Transient overexpression of *AtNPR1* in *N. benthamiana* demonstrated that AtNPR1 co-migrates with the 70kDa band detected in tobacco extracts (Figure S18). In all cases, we observed constitutive accumulation of NPR1 in non-treated tobacco leaf tissue. However, incubation of cut leaf disks in water or SA or agroinfiltration of *N. benthamiana* resulted in a slightly reinforced NPR1 signal (**Figure 7**; Figures S17 and S18). The data are consistent with our previous observation that the *NtNPR1* gene is expressed constitutively in tobacco leaves and in whole seedlings (Maier et al., 2011).

**Figure 7.**
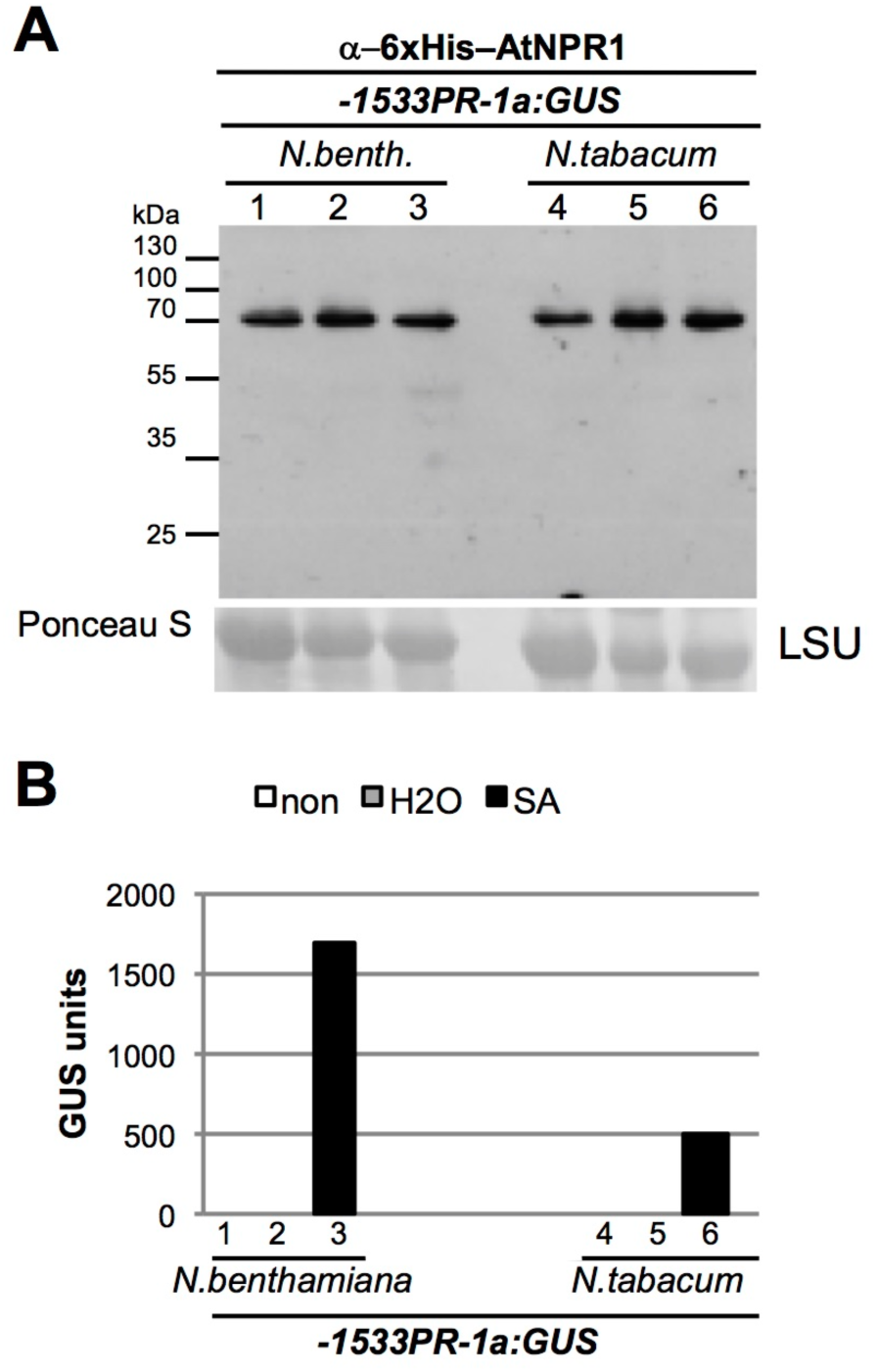
NPR1 accumulates constitutively in tobacco leaves. **(A)** Detection of NPR1 in leaf extracts of *N. benthamiana* and *N. tabacum*. Proteins were extracted from non-treated tissue (1,4), from water-floated leaf disks (2,5) and from SA-floated leaf disks (3,6) and separated by SDS gel electrophoresis. NPR1 was detected with an antiserum raised against 6xHis–AtNPR1. Staining of the large subunit of RUBISCO (LSU) with Ponceau S demonstrates loading of the nitrocellulose filter. **(B)** GUS activities in leaf extracts of *N. benthamiana* and *N. tabacum* carrying the *-1533PR-1a: GUS* reporter gene. The same extracts as used for immunodetection of NPR1 were tested.

## DISCUSSION

In search for components of the NPR1-mediated signaling pathway, we identified TGA7, a novel member of the TGA family of bZIP transcription factors from tobacco. Sequence alignments and functional analyses group TGA7 to clade III. Including TGA7, the tobacco TGA family now contains six members belonging to four clades. *TGA7* cDNA clones were isolated in a Y2H screen with the AtNPR1 bait. Hence, TGA7, like Arabidopsis TGA2 and TGA3, would qualify as transcription factor associated with the pathogen defense response SAR.

### Functional role of TGA7

Indeed, using an artificial reporter gene system, we show that chimeric TGA7GBD functions as transcription factor that is specifically activated by SA in young leaves of tobacco plants. Reporter enzyme activity is very low in unchallenged plants while it increases more than 10-fold in response to SA and its functional analogs. Our findings corroborate previous observations on an *Arabidopsis* TGA2GBD chimera using the same reporter gene system (Fan and Dong, 2002). Most intriguingly, however, we could not detect *TGA7* transcripts in young leaf tissue where SAR normally occurs. In addition, previous results have shown that tobacco clade II factor TGA2.2 is the main component of the ASF-1 complex from tobacco leaves mediating activation of SA-responsive genes, e.g., *PR-1*, in the course of SAR (Niggeweg et al., 2000b). Therefore, although TGA7 clearly possesses the potential to activate gene transcription in response to SA, it remains indefinite whether TGA7 indeed plays a role in the course of SAR.

Yet, we detected *TGA7* transcripts in leaves of older tobacco plants and in whole seedlings. Here, increased expression of the endogenous *TGA7* gene parallelled transcription activity of the TGA7GBD chimera. Together, the data suggest that TGA7 may fulfill a role at these specific developmental stages. TGA7GBD transcription activity in older leaf tissue was coincident with gene expression from the *PR-1a* promoter, indicating that developmental leaf senescence was initiated in plants. PR-1 proteins accumulate in leaves of flowering tobacco plants along a developmental gradient from higher levels in lower older leaves to weaker levels in upper younger leaves (Fraser, 1981; Grüner and Pfitzner, 1994). It has also been shown that the SA signal plays a role in control of gene expression during later stages of the senescence program in tobacco and *Arabidopsis* (Yalpani et al., 1993; Morris et al., 2000), and that activation of NPR1 promotes SA-induced leaf senescence in *Arabidopsis* (Chai et al., 2014). Thus, it appears plausible that TGA7 could mediate SA-responsive gene transcription in older tobacco leaf tissue.

While activation of TGA7GBD may proceed through SA signaling in senescent leaves, we would exclude this option for cotyledons of tobacco seedlings. Histochemical staining demonstrated that TGA7GBD strongly activates the *UAS^Gal4^:GUS* reporter in cotyledons of seedlings raised on MS mediun without addition of SA. This result is clearly different from previous analysis of TGA2GBD in *Arabidopsis* which remained inactive in seedlings cultivated on MS medium (Fan and Dong, 2002). Our findings support the notion that clade II factor AtTGA2 and clade III factor TGA7 may have different functional significance in planta. In fact, we cannot anticipate elevated levels of SA in cotyledons, since SA-responsive promoters of the SAR marker genes *PR-1a*, *NIMIN1* and *NIMIN2* are not active in tobacco seedlings grown on MS medium (Hermann et al., 2013), and PR-1 proteins do not accumulate in seedlings with TGA7GBD-mediated reporter activity. It therefore appears that TGA7 is able to mediate both SA-dependent and SA-independent gene transcription in tobacco leaf tissue, albeit at distinct developmental stages.

In young tobacco leaves, the TGA7GBD chimera remained largely inactive in absence of SA. This finding is noteworthy, as expression of the *TGA7GBD* gene was under control of the strong constitutive *CaMV 35S* promoter. In yeast, TGA7 can interact with other TGA factors. In addition, TGA7 is able to dimerize with the SAR regulator NPR1 and with glutaredoxins. All interactions likewise occur on the conserved C-terminus of TGA7. We do not know the precise binding sites for these different proteins on the TGA7 C-terminal end. However, we would suppose that binding to the TGA7 C-terminus is mutually exclusive and likely linked to distinct functions of different protein complexes formed in planta.

### Role of NtNPR1 in control of TGA7 activity

Regarding NPR1, interaction with TGA factors occurs with N-terminal regions of AtNPR1 and NtNPR1 encompassing the central ankyrin repeat domain. In contrast, TGA7 binds to the conserved C-terminal third of NtNPR1 lacking ankyrin repeats. The NtNPR1 C-terminal region does not harbor general protein-protein interaction motifs, but was shown to display responsiveness to the SA signal, exerted via two domains conserved among NPR family members in higher plants, the LENRV motif and the binding domain for NIMIN1 and NIMIN2 proteins (N1/N2BD; Maier et al., 2011). In yeast, interaction with NPR1 dampens TGA7 transcription potential, indicating that NPR1 can negatively impact TGA7 activity. Expression of *TGA7* and interaction of TGA7 with the NtNPR1 C-terminus are not affected by SA levels. However, interaction of TGA7 with the NtNPR1 C-terminus is affected by presence of NIMIN2-type proteins. We found that NIMIN2 proteins compete with TGA7 for interaction with the NtNPR1 C-terminus, and that TGA7 binds to the NtNPR1 C-terminus in vicinity to the N1/2BD. Thus, accumulation of NIMIN2 proteins in response to SA, occuring during SAR and during developmental leaf senescence (Horvath et al., 1998; Zwicker et al., 2007), could induce relief of TGA7 and chimeric TGA7GBD from NtNPR1, which is present constitutively in tobacco leaves. In this scenario, NtNPR1 would act as repressor of TGA7 and TGA7GBD transcription activity, and SA-induced NIMIN2 proteins would promote transcription through liberation of TGA7 from NPR1. After removal from NPR1, TGA7 could commit itself to novel interactions with other partner proteins, e.g., promoter-bound TGA factors, to elicit TGA7-dependent gene induction in response to the SA stimulus. Alternatively, TGA7, itself, could bind to the promoter regions of SA responsive genes. In this line, the SA receptor NPR3 has been demonstrated to repress the transcriptional defense response during early flower development in *Arabidopsis* (Shi et al., 2013). In fact, gene induction via removal of inhibiting repressor proteins has been recognized as general theme in other plant hormone signaling cascades (Robert-Seilaniantz et al., 2011; Arabidopsis Interactome Mapping Consortium, 2011; Pieterse et al., 2012; Campos et al., 2014). Of note, our findings are not in conflict with the established role of NPR1 as co-activator of *PR-1* gene induction during the SAR response (Rochon et al., 2006; Withers and Dong, 2016). Together, the results suggest that transcription activator TGA7 is controlled in different ways, first at the transcription level by gene expression occurring preferentially at distinct developmental stages and in specific tissues, and second by posttranslational regulation through interaction with various partner proteins, notably NtNPR1, able to modify TGA7 activity.

### A tentative model for SA- and TGA7-mediated gene activation

In our current working model for SA- and TGA7-mediated gene induction in tobacco, TGA7 transactivation potential is modulated by the NtNPR1 C-terminus, thus imposing SA dependency on TGA7 transcription activity (**Figure 8**). In presence of NtNPR1, TGA7 is inactive. Rising levels of SA lead to induction of NIMIN2-type proteins which are able to replace TGA7 from the NtNPR1 C-terminus. Liberated TGA7 can then join other TGA factors to initiate expression of target genes with *as-1*-like cis-acting elements. We suggest the depicted mechanism to occur in the course of developmental leaf senescence in tobacco. We further propose that SA-dependent gene induction during senescence and SA-dependent gene expression during SAR may follow alternative routes, while, on the other hand, both pathways engage the same signal, related signaling components, and likewise result in activation of a similar set of genes, f. e., *PR-1* (Morris et al., 2000). Consistent with previous findings, we suspect that the SAR pathway, besides NPR1, relies on transcription factors of the TGA family other than TGA7 and possibly also on other mechanisms of transcription activation. Thus, gene expression during age-dependent senescence and during SAR would be controlled through separate pathways, uncoupling the normal developmental program during the ontogeny of plants from defense responses induced by environmental cues. Similarly, it was postulated that *PR-1* gene expression during HR and during SAR is induced by distinct signaling pathways (Tsuda et al., 2013) which converge on the *as-1*-like cis-acting element of the *PR-1a* promoter (Grüner et al., 2003).

**Figure 8.**
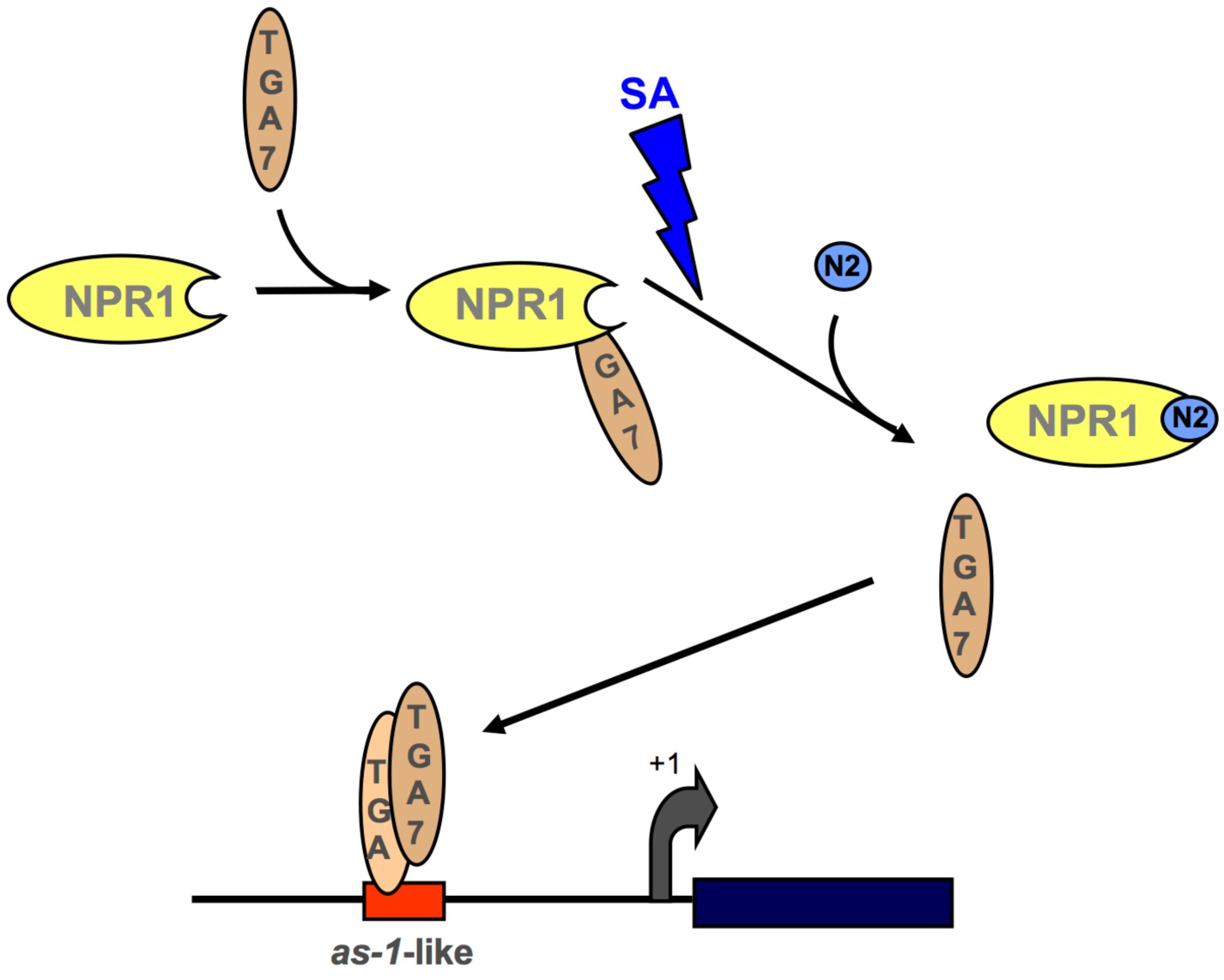
Working model for TGA7-mediated and SA-dependent gene activation in senescent tobacco leaves. In noninduced leaves, TGA7 transcription activity is abrogated by interaction with the NPR1 C-terminal region. Increasing levels of SA in senescent leaf tissue induce accumulation of NIMIN2 (N2) proteins leading to relief of TGA7 from NPR1. Freed TGA7 can hybridize with promoter-bound TGA factors to induce expression of target genes harboring *as-1*-like cis-regulatory elements.

## Conclusions

By characterizing TGA7, a novel member of the TGA family of transcription factors from tobacco, we demonstrate that NtNPR1 possesses two distinct TGA factor interaction sites. Of note, binding of TGA7 to the NtNPR1 C-terminus correlates with low level transcription activity of GBD–NtNPR1 in yeast (Maier et al., 2011). Among members of the NPR family tested by us, i.e., *Arabidopsis* NPR1 through NPR4 and tobacco NPR1 and NIM1-like1, transcriptional activity in yeast is unique to NtNPR1, and thus could be due to interaction with endogenous yeast bZIP proteins on the NtNPR1 C-terminus. Differential interaction with TGA factors enables NtNPR1 to form diverse complexes which may mediate signal transduction through different pathways. Altogether, members of the NPR and TGA factor families and their interaction partners appear to form a highly complex and interconnected network targeting yet unknown effector genes and facilitating coordination of distinct signaling cascades via biochemically related constituents. Finally, our find of TGA7, its binding to the NtNPR1 C-terminus and its displacement by SA-induced NIMIN2 proteins reveal a novel mechanism for SA-dependent gene activation. Our results underline the importance of the NtNPR1 C-terminal region as vital regulatory domain, while they render TGA7 a transcription factor that can be modulated by NPR1, but still permits gene expression independent of the SAR signal SA.

## AUTHOR CONTRIBUTIONS

VS-Z performed Y2H screens and most of the experiments. DN determined protein-protein interaction strengths by Y2H and Y3H assays. EK performed Y2H tests and immunodetection of yeast proteins. MM analyzed gene expression by RT-PCR. MH performed Y1H analyses. FM analyzed the subcellular localization of TGA7–mGFP4 fusion protein in plant cells. VH performed agroinfiltrations. AJPP provided inestimable advice and fruitfull discussions on the project. UMP generated gene constructs, performed immunodetection, was responsible for the coordination and supervision of the work, and wrote the article.

## ACKNOWLEDGMENTS

We are grateful to Nam-Hai Chua (Rockefeller University, USA) and to Xinnian Dong (Duke University, USA) for providing vectors encoding the GBD-based chimeric *GUS* reporter gene system for plants. We also would like to thank Christiane Gatz (Georg-August-Universität Göttingen, Germany) for the generous gift of cDNA clones encoding tobacco TGA factors 2.1, 2.2 and 10 and Eric Lam (Rutgers University, USA) for cDNA clones encoding *Arabidopsis* TGA2 and TGA6; Frederik Börnke (Universität Potsdam, Germany) for the tobacco Y2H cDNA library; Jim Haseloff (University of Cambridge, United Kingdom) for a cDNA clone encoding mGFP4; and Michael Schweizer (University of East Anglia, Norwich, United Kingdom) for providing Y1H vector pRW95-3. Finally, we would like to thank Simone Mast for help with Y2H screening, Sylvia Zwicker for performing Y2H interaction assays, Christoph Bäuscher and Christine Arnold for protein expression and purification and Ingrid Prießnitz-Hohos for plant transformation. This work was supported by Deutsche Forschungsgemeinschaft (DFG; grant nos. Pf 188/8-1 and Pf 188/8-3 to AJPP) and by Universität Hohenheim.

## SUPPLEMENTARY MATERIAL

